# Introducing Haptic Feedback in Real-Time fMRI Neurofeedback: A Novel Approach to Modulate Primary Motor Cortex Activity

**DOI:** 10.1101/2024.09.30.614977

**Authors:** Mathis Fleury, Pauline Cloerec, Quentin Duche, Giulia Lioi, Anatole Lécuyer

## Abstract

As fMRI-Neurofeedback (fMRI-NF) is still in its early stages, many questions remain regarding the optimal methodology, particularly in relation to feedback modalities. One of the core components of neurofeedback is the feedback itself, which the participant relies on to regulate, learn, and refine their mental strategies. However, most fMRI-NF protocols to date have primarily used visual feedback, which may not be ideal in all cases. Certain individuals or populations might benefit from alternative or additional feedback modalities, such as haptic feedback —a novel approach in this field— along with auditory, virtual reality/immersive environments, or a combination of these. In this study, we assess the performance of neurofeedback elicited by a motor imagery (MI) task using visuo-haptic feedback and compare it to unisensory visual and haptic feedback. Our results suggest that combining visual and haptic feedback in neurofeedback may be more engaging than conventional visual feedback alone, particularly in neurorehabilitation, by more effectively activating the primary motor cortex—a region considered a key target for promoting motor recovery.

## 1 INTRODUCTION

Real-time functional magnetic resonance imaging (fMRI) Neurofeedback (NF) is a technique that allows to learn voluntary control over spatially localised brain activity Weiskopf et al. (2004); decharms (2007); Sulzer et al. (2013); Sitaram et al. (2017). It can be used to study causal brain-function relationships by investigating how learned self-regulation of brain activity affects perception or behaviour. Some recent technological advances such as machine learning analyses, wireless and real-time recordings have increased interest in NF and brain-computer interfaces (BCI) approaches, especially electroencephalogram (EEG)-based BCI/NF.

As the fMRI-NF is still in its early days, there are many open questions about the optimal methodology. One of them concerns the feedback modality. Indeed, one of the cornerstones of NF and BCI is the feedback given to the subject whom relies on it to regulate, learn and improve his or her mental strategy. However, to date, most fMRI-NF protocols have only relied on visual feedback Lioi et al. (2018); Mehler et al. (2019), and its use may seem questionable in some cases. As suggested by Stoeckel and colleagues, some people or population might benefit from haptic, auditory, virtual reality/immersion, or the combination of some of these modalities for NF Stoeckel et al. (2014); Paret et al. (2019). Although suggested, few studies have focused on the value of using other feedback modalities. This lack of studies is not apparent in the EEG-NF, where many studies used haptic as feedback modality Buch et al. (2008); Ramos-Murguialday et al. (2012); Fleury et al. (2020).

Even if visual feedback has been shown to be the type of sensory input that produces the best learning processes Hinterberger et al. (2004), there are arguments that would support haptic as a feedback modality. For example, visual feedback may not be suitable for individuals with impaired vision or during a mental motor imagery task, which requires great abstraction from the subject. In this case, a haptic feedback could seem more appropriate and more natural than visual feedback Cincotti et al. (2007). Besides, it has been suggested that providing haptic feedback could improve the sense of agency, a technology acceptance-related factor, in motor imagery (MI) BCI’s Thurlings et al. (2012). Nevertheless, the combination of multiple types of feedback, referred to as multisensory feedback, is expected to provide enriched information Gürkök and Nijholt (2012). However, to be efficient, feedback should not be too complex and should be provided in manageable pieces Lotte et al. (2013). Perceptible gains from the use of different modalities are still little known, no studies have addressed the role of feedback in fMRI-NF.

Applications related to haptic-based BCI are multiple, such as rehabilitation and entertainment. The majority of the clinical papers focus on stroke rehabilitation, because haptic-based BCI/NF seems to be a promising way for motor rehabilitation, as this non-invasive technique may contribute to closing the loop between brain and effect Fleury et al. (2020). The haptic feedback presents 3 main advantages for motor rehabilitation : the production of a kinesthetic illusion, the strengthening of the sensorimotor loop, and a faster action than the visuomotor loop Botzer and Karniel (2013). Moreover, there is frequently a visual handicap (homonymous lateral hemanopsy) linked to the location of the stroke with motor deficit in the upper limb. By providing immediate sensory feedback contingent upon the contraleteral brain activity, we hypothesised that reestablishing contingency between ipsilesional cortical activity related to planned or attempted execution of finger movements and proprioceptive (haptic) feedback, such feedback will strengthen the ipsilesional sensorimotor loop fostering neuroplasticity that facilitates motor recovery.

Haptic systems are categorised into : tactile feedback (vibrotactile, pressure/contact, thermal or electrotactile interfaces); and kinaesthetic feedback, which gives information about the position of the limbs or applied forces (grounded force feedback, exoskeleton devices or functional electrical stimulation). However, many of these systems suffer from problems related to MR-compatibility or are difficult to set up. Vibration stimulation seems to be a good starting point for the creation of MR-based feedback as many technologies allow its introduction into the MR environment (piezoelectric or air pressure devices). Moreover, vibratory stimulation is already used in various medical applications such as pain management or proprioceptive rehabilitation after stroke Murillo et al. (2014); Rittweger (2010). Its role is not only tactile but also proprioceptive because if applied under certain conditions (frequency of 60-100 Hz, tendon target)Gomez-Rodriguez et al. (2011), it can create movement illusions also called kinaesthetic illusions by stimulating the brain motor areas Naito et al. (2002).

The multisensory aspect of feedback seems to be a way to improve the design of NF studies. However, multisensoriality should not just be additive but coherent and synergistic. That is why the association of a virtual hand and the tendinous vibration on the wrist of this same hand seems to be an ecological tool for an MI task. It allows the user to perceive a vibration at the level of his hand (with potential hand movement illusion) while having a visual illusion. In this study, we used tendon vibration associated with a VR environment as a multisensory feedback with a twofold rational: firstly to obtain a potentiation of the illusion, and secondly because of the observation of a motor task playing a role in brain motor activations.

To our knowledge, this is the first fMRI-NF study to incorporate haptic feedback alongside visuo-haptic feedback. The present study aims to advance this field by: (1) evaluating the effectiveness of haptic feedback in motor imagery (MI)-based fMRI-NF, (2) developing a more ecologically valid, multisensory approach by combining immersive visual feedback with haptic feedback, and (3) comparing the performance differences between unisensory (visual or haptic) and multisensory feedback modalities. To achieve this, we used an MR-compatible vibrotactile device for delivering haptic feedback. We hypothesize that our multisensory approach will (1) create a more immersive and engaging feedback environment, (2) enhance participants’ ability to regulate and learn through neurofeedback, and (3) offer a more effective alternative to conventional unisensory feedback methods.

## 2 MATERIALS AND METHODS

### 2.1 Participants

Fifteen healthy adult volunteers were involved in this study (5 women, *meanage* = 27, *SD* = 3.26). All participants met the following inclusion criteria: aged between 18 and 80 years, right-handed, with no history of neurological conditions such as brain injury, brain surgery, or epilepsy. Exclusion criteria included left-handedness and vision impairments beyond -3 to +3 diopters when correction with contact lenses was not feasible.

### 2.2 MRI Data Acquisition

All MR images were acquired from a Siemens 3T scanner (MAGNETOM Prisma, Siemens Healthineeers, Erlangen, Deutschland) with 64-channel head coil. MRI data were acquired on the Neurinfo MRI research facility from the University of Rennes I.

Functional data is obtained with a T2*-weighted single-shot spin-echo EPI with the following parameters Moeller et al. (2010): repetition time (TR) / echo time (TE) = 1000/30 ms, Field-Of-View (FOV) = 210 × 210 mm^2^, 42 slices, voxel size = 2.5×2.5×3mm^3^, matrix size = 105 × 105, flip angle = 65°, multiband acceleration factor 3. As a structural reference for the fMRI analysis, a high resolution 3D T1 MPRAGE sequence was acquired with the following parameters: TR/ inversion time (TI) /TE = 1900/900/2.26 ms, GRAPPA 2, FOV = 256 × 256 mm2 and 176 slabs, voxel size = 1 × 1 × 1 mm^3^, flip angle = 9°.fMRI data were pre-processed online for motion correction, slice-timing correction and fMRI NF features were then computed in the NF control unit using a custom made script developed in Matlab 2017 (The MathWorks, Inc., Natick, Massachussets, United States) and SPM12 .

### 2.3 Experiment Design

The overview experimental design is summarised in *Figure 1*. All participants were engaged in one fMRI-NF session on one day. Each participant performed 3 training blocks, in which they received Visual (NF-V), Haptic (NF-H) and Visuo-Haptic (NF-VH) feedback, using a counterbalanced order across participants who were blinded to the order Keedwell and Dénes (2015). An oral information was given just before the session about MI, NF and movement illusion. Instructions for MI oriented the volunteers towards a kinesthetic MI which involves taking a first-person perspective and imagining the feeling and experience of movements without overt movement Hanakawa (2016). Participants were naive with respect to the purpose of the experiment.

**Figure 1.**
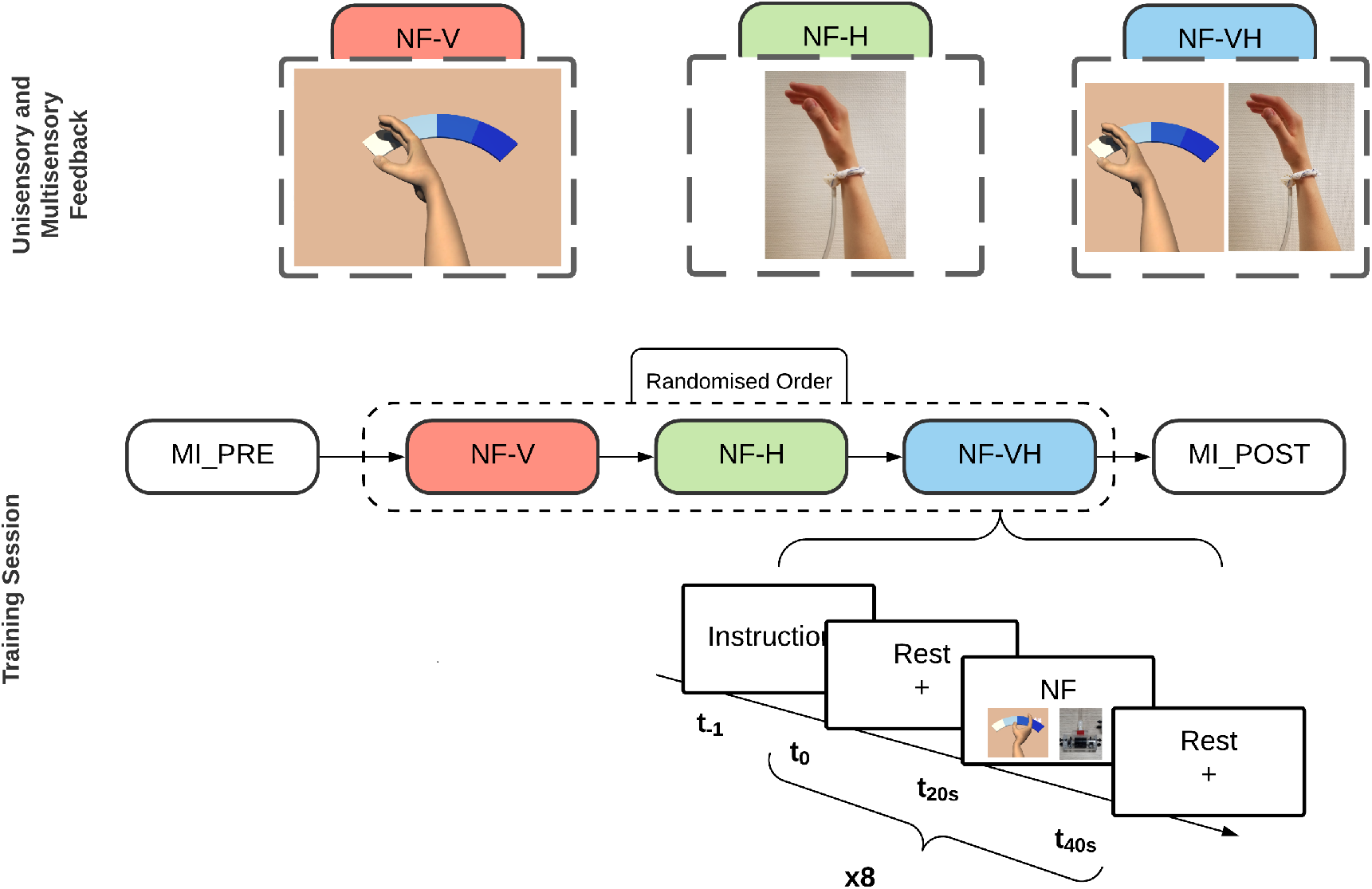
A schematic representation of the experimental protocol is provided. The first row illustrates the types of feedback presented to participants during the training runs: NF-V represents visual feedback, NF-H refers to haptic feedback, and NF-VH denotes multisensory feedback combining both visual and haptic inputs. The second row outlines the training sessions, where it is important to note that the three types of feedback are randomized for each participant using a Latin square design.

A pneumatic vibrator was set on the flexor carpi tendon of the right hand (dominant hand) at the beginning of the session when the volunteer was lying in the MRI. It was maintained on skin with velcro strip (Figure 1 *NF-H*). An important point is that the hand was in a resting position, and did not touch anything around.

The session consisted of five MI functional runs and an anatomical scan (T1 MPRAGE). Each functional run consisted of 8 blocks of rest (20 s) and tasks (20 s). During the 1st run, the participant did not receive any feedback (first no-feedback) but only instructions: *“Imagine moving your right hand”* for the task and a dark grey cross for the rest. During the runs 2 to 4, the various feedback on the activation level of the selected target region were given (feedback runs). After the training runs, participants were engaged in one last run in which again no feedback was given (last no-feedback). Randomised condition orders were equally balanced over all sessions of all participants.

At the end, each participant filled a post hoc questionnaire, gathering qualitative information about the different feedback. In the questionnaire, subjects had to express their degree of agreement about each affirmation by using a Likert-scale from 1 to 6 (1 = totally disagree, 6 totally agree).

### 2.4 Region-of-Interest (ROI) and Calibration

For the fMRI calibration and the definition of ROIs, data of the motor imagery session (no-feedback) were pre-processed for motion correction, slice-time correction, spatial realignment with the structural scan and spatial smoothing (6 mm FHWM Gaussian kernel). A first-level general linear model (GLM) analysis was then performed. The corresponding activation map was used to define two ROIs around the maximum of activation in left M1 and left SMA. To this end, two large apriori masks were defined and the respective ROIs identified taking a box of 9 x 9 x 3 voxels (20 x 20 x 12 mm^3^) centered around the peak of activation (thresholded T-map *task > rest, p <* 0.001, *k >* 10) inside the apriori masks. The position of the ROIs was validated by a clinician. A weighted sum of the BOLD activity in the two ROIs was then used to compute the fMRI NF (Section 2.5). Also for the fMRI NF, a threshold was set by estimating the value reached 30% of the time during the calibration session.

### 2.5 Real-time fMRI system and NF calculation

Real-time neurofeedback (NF) calculations, detailed in Perronnet et al. (2020), were carried out on a dedicated computer (Intel Core i7, 16 GB RAM, Windows 10). The fMRI NF signal was computed as the difference in percentage signal changes between two regions of interest (ROIs)—the supplementary motor area (SMA) and the primary motor cortex (M1)—and a large deep background region (slice 3 of 42) with activity unrelated to the NF task. This approach was used to minimize the impact of global signal changes caused by factors such as breathing, heart rate, and head movements ( Thibault et al. (2018)). The NF feature was calculated using the following equation:

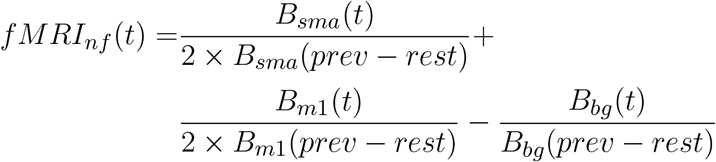

*B*_*sma*_ is the average bold signal in the SMA ROI, *B*_*m*1_ in the M1 ROI and *B*_*bg*_ in the background slice. *B*_*x*_(*prev* − *rest*) is the ROI *x* baseline obtained by averaging the signal in the ROI *x* from the fourteenth to the nineteenth second (to account for the hemodynamic delay) of the previous rest block. The same weight is given to both ROIs. The fMRI feature was smoothed over the three last volumes, divided by the individual threshold and eventually translated as feedback every repetition time (1 s).

### 2.6 Set-up: Unisensory and Multisensory feedback

#### 2.6.1 Visual Feedback

Visual feedback of the performed mental task was given to the participants by using Unity software (version 3.5), and the virtual scene was composed of a homemade neutral and white skin upper limb avatar (cf. 1 *NF-V*). The feedback was a right hand rendered from the point of view of the virtual avatar and moving along a coloured band on a blue scale. The movement executed by the virtual hand was an extension of the right wrist which is congruent with the illusion of movement caused by the haptic feedback Naito et al. (1999).

#### 2.6.2 Haptic Feedback

##### MR-compatible Vibrator

The haptic feedback is a tactile interface based on vibrotactile stimulation. Vibrations are delivered through a pneumatic vibrator which is MR-compatible. The body of the vibrator is a cylinder made of non-magnetic materials, and it contains a wind turbine to which an off-centered mass is attached. The rotation of the off-centered mass generates tangential vibrations transmitted to the vibrator body. The vibration frequency and amplitude depend on the angular velocity of the rotor, which is proportional to the air inflow. The device is controlled through a system placed outside the scanning room. The maximum frequency intensity of a pneumatic vibrator is dependent on the input air pressure. In our case, the system was capped at 4 bar, which allows a maximum frequency of 60Hz. This frequency can elicit movement illusion Le Franc et al. (2020, 2021) which begin from 60Hz to 100Hz.

##### Semi-continuous feedback

The vibration frequency of the pneumatic vibrator was used as feedback, frequencies were allocated to map the whole range of NF scores. The vibration was delivered continuously and in order to ensure that the user could perceive the frequency changes. The frequency was selected according to the *just noticeable difference* of the vibrotactile perception ( 20% between each frequencies) Pongrac (2008). Four frequency steps were then allocated as follows:

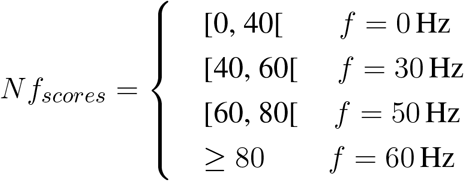

In order to avoid getting occurrences of BOLD reactions in motor cortex and to reward MI task above all. The vibration frequency is 0Hz up to 40% NF scores. The interest is that the subject must be fully engaged in the task before receiving the feedback.

#### 2.6.3 Visuo-Haptic feedback

The visuo-haptic feedback is the combination of the visual and haptic feedback respectively. Visual feedback being a representation of the illusion of movement induced by haptic feedback. Hence, if the virtual hand moves towards the dark blue area, the vibration will be greater and thus the illusion of movement will be intense.

### 2.7 Offline Data Analysis

#### 2.7.1 Functional MRI Preprocessing

Structural and functional MRI data were pre-processed and analysed with AutoMRI, a proprietary software using SPM12 and Matlab. The structural 3D T1 images was segmented into tissue class images (grey and white matter, cerebrospinal fluid compartments, soft tissue, bone and others) and normalised to the Montreal Neurological Institute (MNI) template space. The preprocessing of fMRI included successively a slice-timing correction, a motion correction, a coregistration to the 3D T1 and a spatial normalisation to the MNI space, followed by a spatial smoothing with a 6-mm FWHM Gaussian kernel.

#### 2.7.2 First-Level Analysis

For the first-level GLM analysis, the regulation blocks were modeled as boxcar functions convolved with the canonical hemodynamic function of SPM12. In the GLM, the six parameters of movement (translation and rotation) were included as covariates of no interest. For each task of each participant, a positive contrast between NF regulation and baseline (rest) blocks was applied.

#### 2.7.3 Whole-brain Analysis

Contrast images were thresholded at *p <* 0.001 and false discovery rate (FDR)-corrected on cluster level at *p <* 0.05. Group-level images were visualised in a sliced brain using Nilearn (http://nilearn.github.io/).

#### 2.7.4 Offline ROI Analysis

We performed offline analysis on regions of interest (ROIs) defined using binary masks derived from the Human Motor Area Template (HMAT) atlas Mayka et al. (2006). The HMAT atlas was utilized to extract six specific motor regions: primary motor cortex (M1), supplementary motor area (SMA), pre-supplementary motor area (preSMA), ventral premotor cortex (PMv), dorsal premotor cortex (PMd), and primary somatosensory cortex (S1) from preprocessed BOLD images.

For each defined ROI, we calculated the mean t-score across various contrasts obtained from task-related fMRI data. Subsequently, we performed an analysis of variance (ANOVA) to investigate the effects of task-related fMRI contrasts within M1 and SMA. Following the ANOVA, we employed Tukey’s Honestly Significant Difference (HSD) test to evaluate the statistical significance of differences between specific contrasts. All parametric statistical analyses were conducted using Python 3.11 (and the statsmodels library).

#### 2.7.5 Questionnaires

All data were analysed by statistical tests using Jamoji 1.1.9.0 and RStudio. Qualitative variables are represented with numbers and percentages, and quantitative variables are represented with means (standard deviations). A within-group analysis comparing the 3 conditions have been performed using Friedman tests (the non-parametric approach was used because N is relatively small).

## 3 RESULTS

The following section shows a comparative analysis between visual, haptic and visuo-haptic session in terms of NF performance and fMRI ROI analysis.

### 3.1 NF Performance

In the offline data analysis, we found a significant difference (*p <* 0.001) in activation in the left M1 ROI between rest and task by studying all runs and all subjects together.

Concerning the NF score of M1 and SMA during the three NF runs, the activity within SMA during the NF-H and NF-VH runs was significantly higher than NF-V (*p <* 9, 58*e* − 10, Kruskal-Wallis test) but no difference was found between NF-H and NF-VH (*p <* 0, 99, Kruskal-Wallis test). The activity within M1 during NF-VH was significantly higher than NF-V (*p <* 0, 05, Kruskal-Wallis test) and NF-H (*p <* 1, 72*e* − 07, Kruskal-Wallis test), NF-V is also significantly higher than NF-H (*p <* 0.004, Kruskal-Wallis test).

### 3.2 Whole Brain Analysis

A whole-brain analysis of the three training sessions—visual (V), haptic (H), and combined visual-haptic (VH)—revealed significant neural activations during neurofeedback (NF) (see Figure 2 A, B, C). Specifically, activation was observed in the pre-supplementary motor area (pre-SMA). When examining the group activation maps for the training runs at an uncorrected significance level of p=0.001p=0.001, common activations during the task were identified (Figure 2 A, B, C).

**Figure 2.**
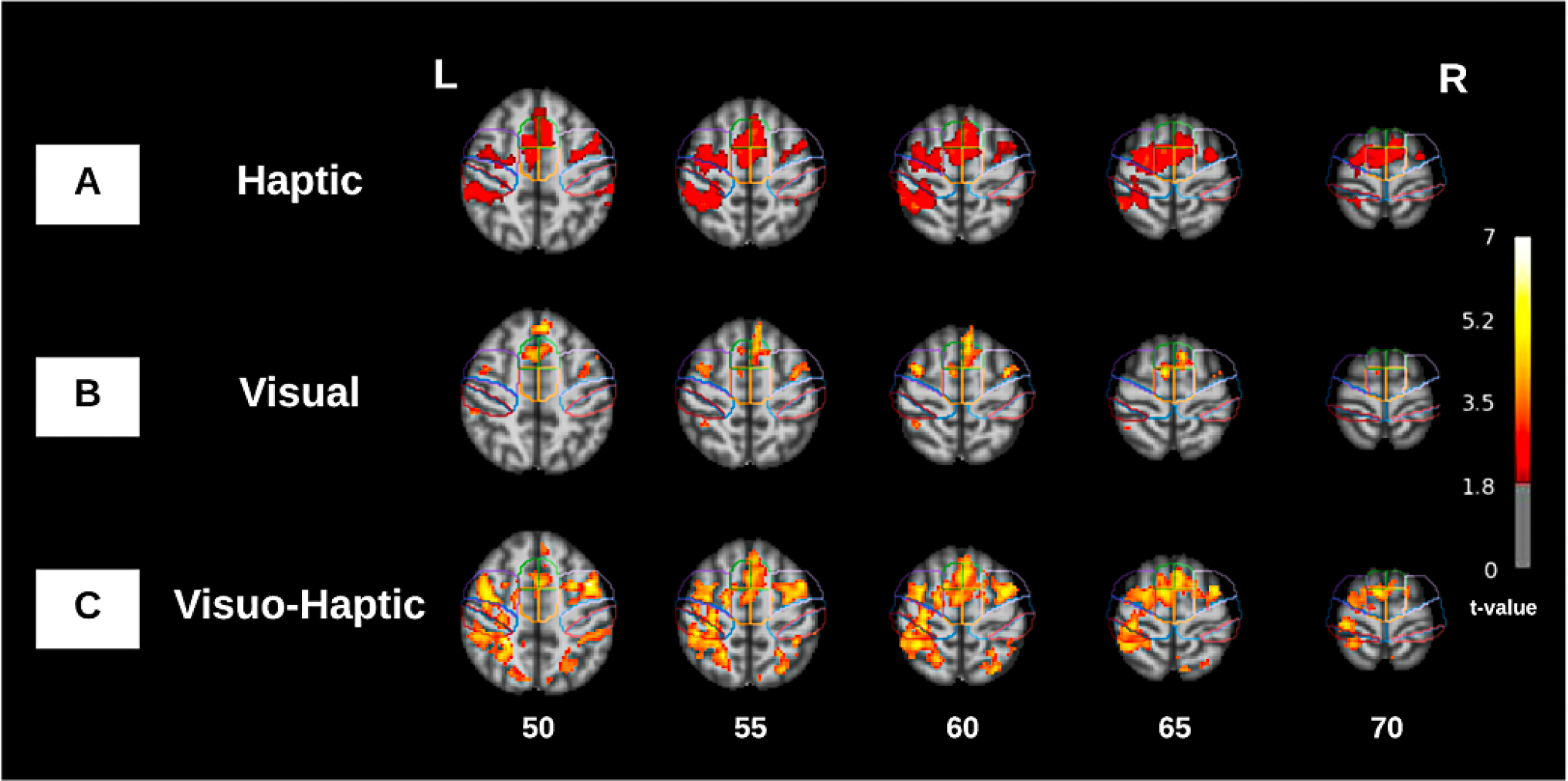
Group activation maps of the training runs in MNI coordinates (*p <* 0.001, uncorrected). The outline of the motor areas of interest based on the HMAT atlas is indicated: preSMA (green) SMA (orange), PMC (purple), M1 (blue) and Sensory motor cortex (red).

In the NF-H run, notable activations were detected in the bilateral pre-SMA and supplementary motor area (SMA), as well as in the bilateral premotor cortex (PMC), the left primary motor cortex (M1), and the left sensorimotor cortex, with T-values ranging from 1.8 to 3.5. Conversely, during the NF-V run, significant activations were found in the bilateral pre-SMA, bilateral PMC, and the left SMA. Although the extent of activation was more limited in this run compared to the haptic feedback condition, the T-values were higher, ranging from 3.5 to 5.2 (Figure 2 B).

In the NF-VH run, extensive activation was again observed in the bilateral pre-SMA and SMA (with a left-side predominance in SMA), as well as in the bilateral PMC and the left M1 and sensorimotor cortex. The significant activation in this condition was broader, with T-values ranging from 3.5 to 5.2 (Figure 2 C).

Regarding contrasts, the comparison between H and V did not yield any significant activations or deactivations (Figure 3 A). However, the contrast between VH and V revealed significant activations in the left M1 (Figure 3 B). Additionally, the comparison between VH and H showed activations within the visual network (Figure 3 C).

**Figure 3.**
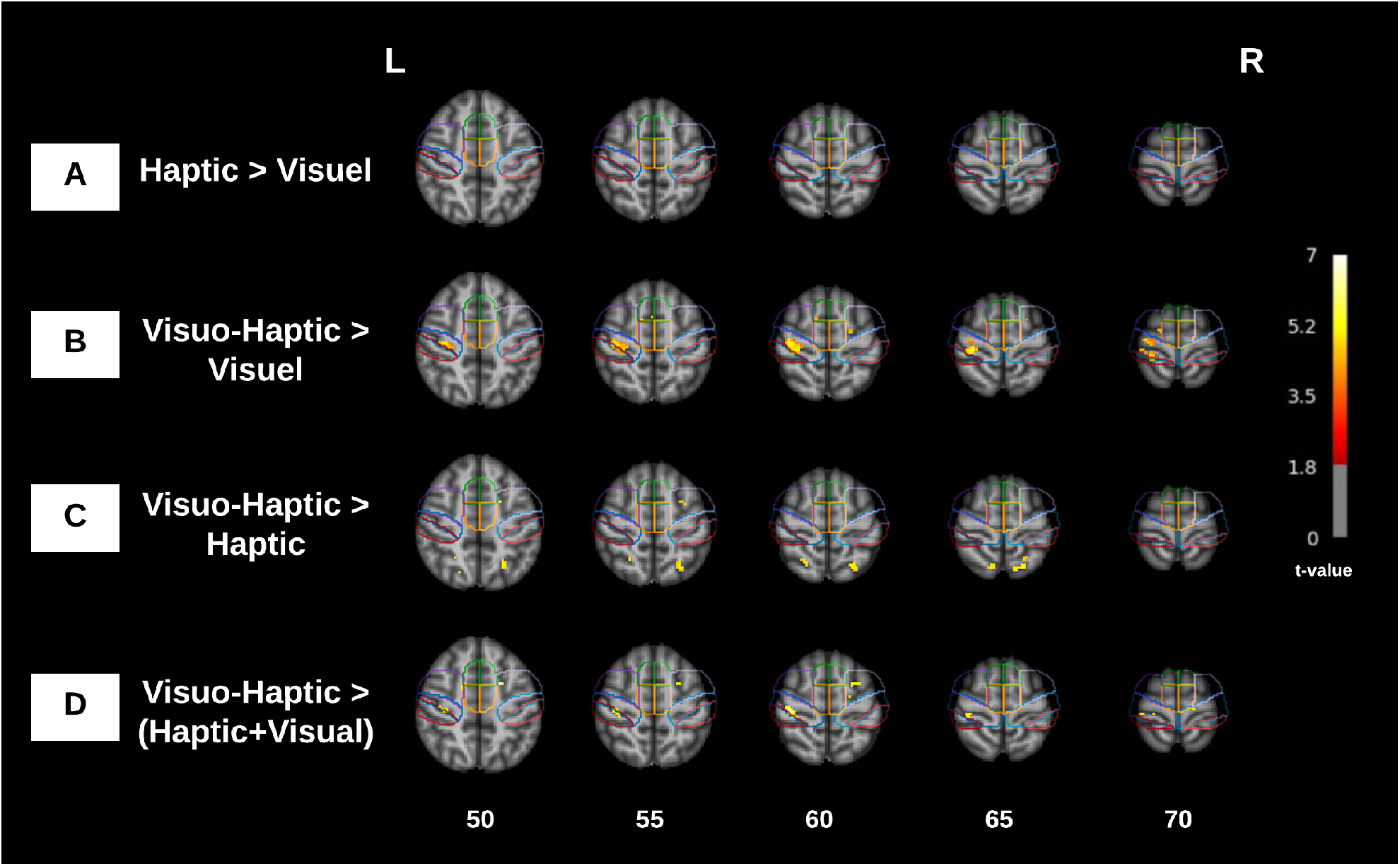
Group activation maps of the training runs in MNI coordinates (*p <* 0.001, uncorrected). The outline of the motor areas of interest based on the HMAT atlas is indicated: preSMA (green) SMA (orange), PMC (purple), M1 (blue) and Sensory motor cortex (red)

Focusing on the primary motor cortex (M1) and supplementary motor area (SMA) as key regions of interest for rehabilitation, we observed activation in the bilateral SMA across all three training sessions (NF-H, NF-V, and NF-VH). Notably, significant activation in the left M1 was only detected during the NF-VH run (Figure 2). When comparing the different feedback conditions using contrasts to assess the differences in activation between the runs, we found a significant activation in the left M1 for the contrast between VH and V.

### 3.3 ROI analysis

An analysis of variance (ANOVA) was conducted to examine the effects of region of interest (ROI) and contrast, as well as their interaction, on the dependent variable. The results revealed that the main effect of ROI was not statistically significant, F(1,84)=0.98, p=0.325, F(1,84)=0.98, p=0.325. Conversely, the main effect of contrast approached statistical significance, F(2,84)=3.15, p=0.048, F(2,84)=3.15, p=0.048, indicating that there are significant differences among the contrasts. However, the interaction between ROI and contrast was also not significant, F(2,84)=0.31, p=0.733, F(2,84)=0.31, p=0.733. Post-hoc analysis using Tukey’s HSD test revealed a significant difference between NF-V and NF-VH (p=0.0008) (See Figure 4), while no significant differences were found between NF-H and the other contrasts.

**Figure 4.**
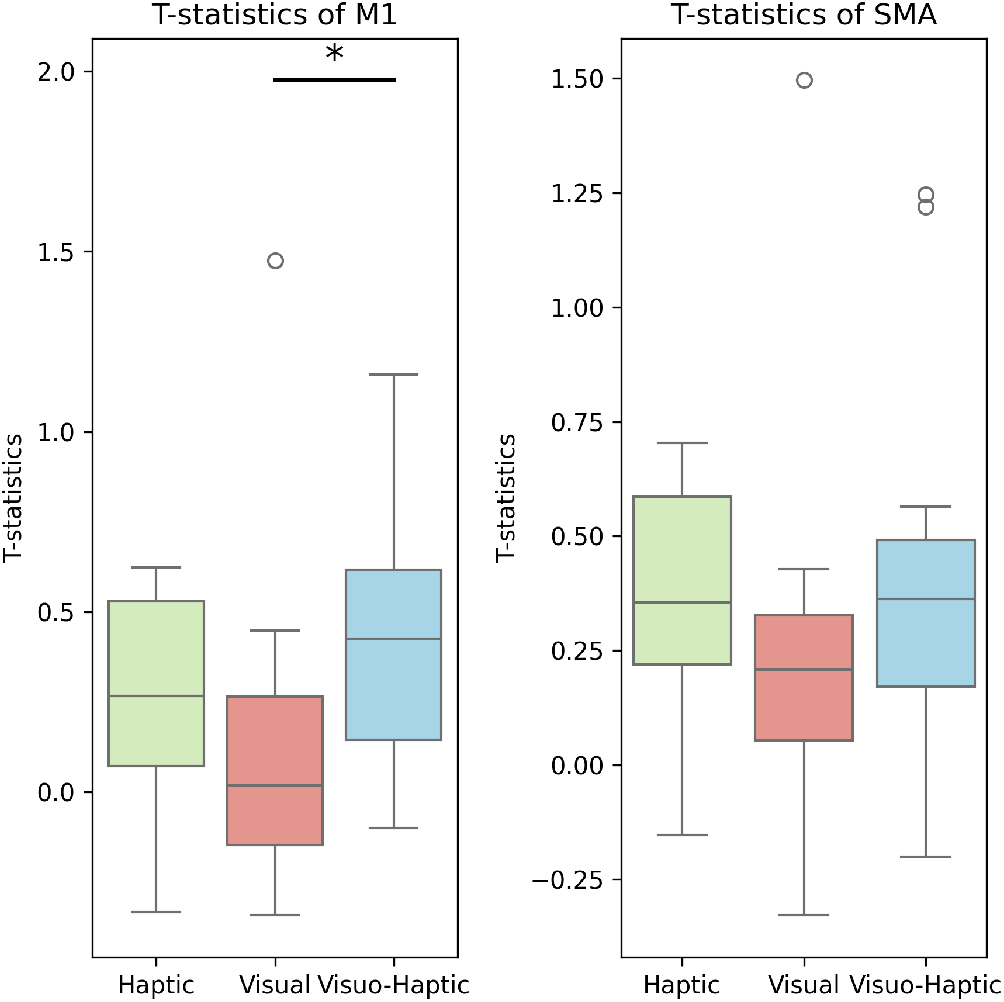
Boxplots illustrate mean differences in t-values between conditions (‘Haptic’, ‘Visual’, ‘Visuo-Haptic’) across regions of interest (SMA and M1). Asterisks (*) indicate statistically significant differences (p *<* 0.05) between conditions.

### 3.4 Mental strategies underlying self-regulation and questionnaires

Among 15 participants, 12 were naive about MI (80 %). We reported the different mental strategies underlying self-regulation : 10 reported having performed a kinesthetic MI (66,7 %). Some reported strategies were opening a door lock with a key, tapping, rotating flexing or extending the wrist.

Concerning the visual feedback, the participants reported a weak appropriation of the virtual hand (Mean = 2.64, DS = 2). The frequently proposed modification for better appropriation was a gesture modification of the hand.

Concerning the haptic feedback, the participants did not report any discomfort with the vibrator. Especially on the affirmation *“the sensation of vibration becomes uncomfortable”*, they did not agree with a mean (SD) likert ranking at 1,53 (1.41). Some participants reported however a transient feeling of paresthesia on their wrist and/or their hand, without any need to stop the experiment.

Concerning the visuo-haptic feedback, they did not agree with the affirmation *“I found the association of two information too difficult to integrate”* with a mean (SD) likert ranking at 2.53 (1.77). Subjects reported to have paid more attention to haptic feedback for 33.3% (N=5), to visual feedback for 13.3% (N=2) and for visuo-haptic feedback for 53.3% (N=8). The degree of agreement about the affirmation *“I found the multisensorial feedback more natural than unisensory feedback”* was equal to a mean (SD) likert ranking at 3.87 (1.60).

Finally, there was no statistical difference between the 3 feedbacks to perform in MI (x2 = 1.32 p =0.517) or concerning the feedback’s reliability during the MI (x2 = 2.21 p =0.074). But to improve the performance of the MI in further experimentation, the most useful feedback among the subjects would be VH for N=9 (60), H for N= 5 (33.3), and V for N = 1 (3.7).

## 4 DISCUSSION

In this study we investigated the use of a multisensory feedback based on visual and haptic as a semi-continuous feedback for MI-NF. We studied its contribution in terms of NF performance and ROI analysis. In order to obtain this multisensory feedback that is congruent with the MI-task, the kinaesthetic feedback was delivered to the subjects coupled with the visual feedback of a virtual arm. The novelty of this approach lies in the creation of continuous MR-compatible haptic feedback, that is provided accordingly to the subject’s MI performance.

The choice of the region-of-interest (ROI) is quite important from the perspective of motor rehabilitation. Kinaesthetic motor imagery seems to be able to activate the same neural networks as real movements in functional imagery Chholak et al. (2019). While the primary motor cortex (M1), which directly controls the execution of the movement, has been suggested to be the most promising target for an efficient motor recovery Favre et al. (2014), supplementary motor area (SMA), which coordinates and plans the movement, seems to be easier to engage during motor imagery Mehler et al. (2019) and more robust than M1.

Subjectively, participants tend to the multisensory feedback : 60 % will choose the multisensory feedback to improve their performance in MI-NF in further experimentation. The NF scores and ROI analysis support this trend to the multisensory. We found a significant higher activation in left M1 on the contrast VH-V.

In a future study it would be interesting to increase the power of vibration in order to obtain more intense illusions of movements, because technical limitations only allowed us to deliver a frequency of 60Hz, which is barely sufficient to deliver an illusion of movement, hence the fact that few subjects (*N* = 2) reported having one.

It could be interesting to test in further studies the same hypothesis with stroke patients to see if similar results could be obtained. This population is often older Lecoffre et al. (2017) than the healthy participants included in our study. We know that the elderly have more difficulties to perceive illusion of movement induced by tendon vibration Chancel et al. (2018) and we can expect a weaker illusion of movement felt with stroke patients because some of them will additionally present cognitive troubles such as attentional ones or sensitive disorders. The current literature Brown et al. (2018); Edwards et al. (2019) remains unclear about the effectiveness of central integration of peripheral vibrations in this population.

## 5 CONCLUSION

In this study, we evaluated the performance of a visuo-haptic feedback for fMRI-NF with a MI task on 15 participants. We compared three conditions: visual alone, haptic alone or the combination of both. The haptic feedback is delivered through a pneumatic vibrator which is MR-compatible. For the visual feedback we used a virtual hand, the movement executed by the virtual hand was an extension of the right wrist. We then compared the BOLD activations as well as the NF scores for the three conditions. The results showed that a visuo-haptic feedback could enable more intense activation of motor regions rather than visual or haptic alone.

## 6 ACKNOWLEDGEMENT

Neurinfo is supported by the the European Union (FEDER), the French State, the Brittany Council, Rennes Metropole, Inria, Inserm and the University Hospital of Rennes.

## Notes

### Competing Interest Statement

The authors have declared no competing interest.

